# Mechanisms Of Meristem Maintenance By Maize Transcriptional Corepressors

**DOI:** 10.1101/2024.04.26.591374

**Authors:** Jason Gregory, Xue Liu, Zongliang Chen, Cecilia Gallardo, Jason Punskovsky, Gabriel Koslow, Mary Galli, Andrea Gallavotti

## Abstract

The formation of the plant body proceeds in a sequential post-embryonic manner through the action of meristems. Tightly coordinated meristem regulation is required for development and reproductive success, eventually determining yield in crop species. In maize, the REL2 family of transcriptional corepressors includes four members, REL2, RELK1 (REL2-LIKE1), RELK2, and RELK3. In a screen for *rel2* enhancers, we identified shorter double mutants with enlarged female inflorescence meristems (IMs) carrying mutations in *RELK1*. Expression and genetic analysis indicate that *REL2* and *RELK1* cooperatively regulate female IM development by controlling genes involved in redox balance, hormone homeostasis, and differentiation, ultimately tipping the meristem toward an environment favorable to expanded expression of the *ZmWUSCHEL1* gene, a key stem-cell promoting transcription factor. We further demonstrate that *RELK* genes have partially redundant yet diverse functions in the maintenance of various meristem types during development. By exploiting subtle increases in ear IM size in *rel2* heterozygous plants, we also show that extra rows of kernels are formed across a diverse set of F1 hybrids. Our findings reveal that the REL2 family maintains development from embryonic initiation to reproductive growth and can potentially be harnessed for increasing seed yield in a major crop species.

**One sentence summary:** REL2-RELKs fine tune hormone and chemical cues to prevent expanded expression of ZmWUSCHEL1 in maize inflorescence meristems, and can potentially be harnessed for increasing seed yield in hybrids.

## INTRODUCTION

The meristem is an organized structure containing a pool of stem cells, which is the product of evolutionary innovation by plants to carefully coordinate multicellular growth during the early colonization of land (NIKLAS AND TIFFNEY 2023). The formation of the shoot system by the shoot apical meristem (SAM) occurs through the repetitive production of leaf and bud primordia, resulting in a succession of phytomers. To achieve this, the meristem is organized into distinct functional zones. The central zone contains the stem cell niche; in the peripheral zone a combination of anticlinal and periclinal divisions ensures a balance of daughter cells that either remain in the niche or enter differentiation into lateral primordia; cells in the rib zone provide the initials for the differentiating ground and vascular tissues of the stem. Sandwiched between the central zone and the rib zone is the organizing center (OC), which contains a small population of cells that express the stem-cell promoting transcription factor WUSCHEL (WUS) (MAYER *et al*. 1998). Our understanding of the molecular mechanisms involved in meristem regulation began with the elucidation of the CLAVATA (CLV)-WUS pathway in Arabidopsis (SCHOOF *et al*. 2000; KITAGAWA AND JACKSON 2019). This negative feedback loop maintains a stable number of stem cells in the central zone ensuring a balance between stem cells and differentiating cells for proper growth. The CLV-WUS pathway involves the transcription of *WUS* in the OC from which it non-cell autonomously activates the central zone specific expression of the *CLAVATA3* (*CLV3*) gene, whose product is processed into a short peptide that is perceived by LEUCINE RICH REPEAT (LRR) RECEPTOR-LIKE (LRR-RLP) and LRR-RECEPTOR-LIKE KINASE (RLK) proteins such as CLV1 in OC cells to repress *WUS* expression (LENHARD AND LAUX 2003) and prevent stem cell over-proliferation in the central zone of the meristem (GUO *et al*. 2010).

In maize, orthologous components of the CLV signaling pathway such as THICK TASSEL DWARF1 (TD1) (BOMMERT *et al*. 2005), and FASCIATED EAR2 (FEA2) (TAGUCHI-SHIOBARA *et al*. 2001) have been identified. The maize genome contains two co-orthologs of *WUS*, *ZmWUS1* and *ZmWUS2* (NARDMANN AND WERR 2006), and recent characterization of the *Bif3* mutant (CHEN *et al*. 2021) demonstrated that ZmWUS1 promotes stem cell fate in the female IM. FEA2 forms two distinct receptor complexes capable of perceiving two distinct peptides: when complexed with the G protein signaling component COMPACT PLANT2 (CT2), FEA2 can perceive the CLV3 ortholog CLE7; alternatively, when complexed with the pseudokinase CORYNE (CRN), FEA2 perceive the differentiating primordia derived ligand FON2-LIKE CLE PROTEIN1 (FCP1) (JE *et al*. 2018). FCP1 can also repress *ZmWUS1* by the receptor FASCIATED EAR3 (FEA3), preventing its expression in the rib zone (JE *et al*. 2016). In a parallel signaling pathway, the bZIP transcription factor FASCIATED EAR4 (FEA4) promotes differentiation in the peripheral zone by positively regulating genes controlling axillary meristem identity, determinacy, and auxin signaling (PAUTLER *et al*. 2015). In line with this, auxin operates antagonistically to cytokinin in the meristem, and promotes differentiation (GALLI *et al*. 2015). As a negative regulator of inflorescence meristem development, the ratio of FEA4’s less active monomeric versus active dimeric state is controlled by the redox environment. Loss of function in OXIDOREDUCTASEs and GLUTAREDOXINs (GRXs) result in an enrichment of oxidized, dimeric FEA4 in the meristem leading to stronger repressive activity and smaller ears (YANG *et al*. 2021).

Similarly, in Arabidopsis the reactive oxygen species (ROS) superoxide (O2^-^) is enriched in the stem cells of the central zone and positively affects *WUS* expression, whereas hydrogen peroxide (H2O2) is enriched in the peripheral zone to promote differentiation. Shifting the O2^-^:H2O2 balance in the SAM alters *WUS* expression, ultimately increasing or decreasing SAM size (ZENG *et al*. 2017). Given the breadth in regulatory control of the meristem, chemical and receptor complex mediated downstream signaling may represent only a portion of the mechanisms employed by plants to ensure proper homeostasis of various meristems.

Mutants of the maize transcriptional corepressor REL2 display pleiotropic vegetative and reproductive phenotypes such as defective axillary meristem (AM) initiation and IM maintenance (GALLAVOTTI *et al*. 2010; LIU *et al*. 2019). Transcriptional corepressors function as molecular bridges between transcription factors (TFs), and MEDIATOR subunits and histone modifiers to silence transcription of downstream target genes (LEYDON *et al*. 2021). REL2 is a functional homolog of the Arabidopsis TOPLESS (TPL) protein, an essential regulator of development (LONG *et al*. 2002; LONG *et al*. 2006; SZEMENYEI *et al*. 2008; SMITH AND LONG 2010). TPL is recruited by WUS in the SAM to repress cell differentiation promoting genes; this interaction is essential for WUS function and lack of it fails to rescue the premature meristem termination of *wus* mutants (KIEFFER *et al*. 2006; CAUSIER *et al*. 2012). However, loss-of-function mutations in *REL2* and its rice ortholog *ASP1* cause enlarged inflorescence meristems (YOSHIDA *et al*. 2012; LIU *et al*. 2019; SUZUKI *et al*. 2019a), an opposite phenotype to *wus* loss-of-function Arabidopsis mutants. How transcriptional corepressors mediate meristem size regulation in monocots is therefore unknown.

## RESULTS

### Isolation and characterization of a genetic enhancer of the *rel2* mutant

To identify genetic modifiers of the *rel2* mutant phenotype, we carried out an EMS mutagenesis screen of the *rel2-ref* allele in the permissive A619 background (LIU *et al*. 2019; LIU *et al*. 2021). We identified two M2 families (M2-01-826 and M2-01-913) that were segregating recessive mutants showing an identical phenotype, namely plants with shorter stature bearing tassels with upright tassel branches (Fig. 1a,b). We provisionally called these mutants *small upright-826* (*sup-826*) and *sup-913*. Crosses between *sup-826* and *sup-913* failed to complement the mutant phenotype in the F1 generation, indicating that the two loci carried different mutations in the same gene. We determined that the small upright phenotype was segregating with a ratio of 1:16 in F2 mapping populations (*sup-826*, 14 mutants out of 222 plants; χ^2^ p=0.95-0.99), indicating that the modifier locus was a true enhancer and did not show an obvious phenotype in a non-sensitized background. We carefully analyzed the phenotype after a few rounds of back-crosses to the original A619 background to remove unlinked EMS generated changes. Double mutant plants were characterized by a severe reduction in overall height (∼40% reduction; Supplemental Fig.1). The reduction in stature was due mainly to a large decrease in internode length, which were approximately half the size when compared to wild type controls, as well as to a reduced number of internodes (on average 1.7 fewer internodes; Supplemental Fig. 1). Inflorescence development was also affected: double mutant tassels tended to be smaller in size (22.5cm vs. 30cm) with a similar number of branches to control plants, while ears were formed on the lowest nodes (1^st^ or 2^nd^ node from the brace root node; Fig. 1a; Supplemental Fig. 1). Approximately 50% of the double mutant plants did not produce an ear (12/25 plants), a phenotype reminiscent of the *rel2* phenotype in the B73 and other genetic backgrounds (LIU *et al*. 2019).

**Fig. 1.**
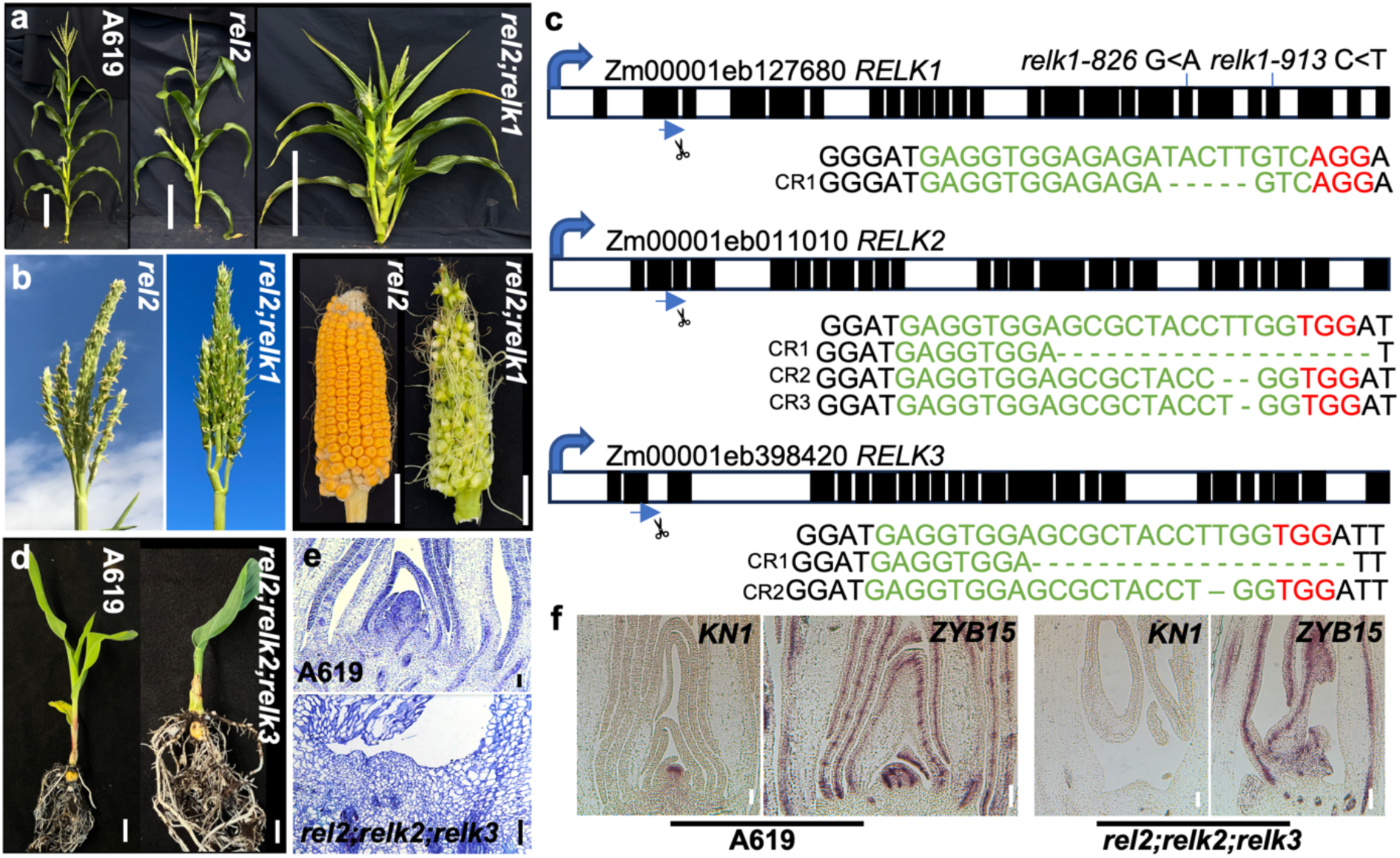
Mutants of *REL2-RELK* are defective in vegetative and reproductive development. (**a**) Representative whole plant images of A619, *rel2,* and *rel2;relk1* plants. Scale bars = 30 cm. *rel2;relk1* plants are smaller, with more upright tassel branches than *rel2* (**b**). Ears produce few viable seeds (**b**). Scale bars = 2.5 cm. (**c**) Schematic representation of the *RELK1* gene with two independent EMS alleles, *relk1-826* in exon 19 and *relk1-913* in exon 22, and one CRISPR-Cas9 generated allele using a single gRNA (green; PAM in red) targeted to exon 2. Alleles of *RELK2* and *RELK3* were generated using a dual targeting gRNA in exon 2. Exons in black, introns in white. (**d**) By 21 days, *rel2;relk2;relk3* has only formed a single visible leaf before apical activity terminates, whereas A619 is forming new leaves. Scale bar = 1 cm. (**e**) Loss of apical activity corresponds to the presence of a SAM-like remnant in seedling sections stained with toluidine-blue (scale bar = 100μm). (**f**) Triple mutants lack detectable *KNOTTED1* (*KN1*) (scale bar = 100μm) but not *ZYB15* (scale bars = 200μm) expression in RNA in situ hybridizations. Wild type sections (**f**) display a normal SAM (scale bar = 100μm), with *KN1* and *ZYB15* expression. n = 2-3 per genotype per experiment.

### The *small upright* phenotype is caused by mutations in the transcriptional corepressor RELK1

Double mutant *sup* plants were crossed to the inbred line B73 to create F2 mapping populations for positional cloning purposes (GALLAVOTTI AND WHIPPLE 2015). We performed whole genome sequencing on the double mutant bulks (14 samples for *sup-826* and 18 samples for *sup-913*). In both cases, the mutant mapped on chromosome 3 and SNP analysis identified in each mutant bulk sample a base change in the coding sequence of gene model *Zm00001eb127680* (B73v5), causing premature stop codons in amino acids W792 and Q939 in *sup-826* (transition G to A) and *sup-913* (transition C to T), respectively (Fig. 1c). *Zm00001eb127680* corresponds to *RELK1*, another member of the *REL2* family of transcriptional corepressor (LIU *et al*. 2019). The corepressor proteins of the REL2 family are homologs of the Arabidopsis TPL/TPR family and contain conserved protein-protein interaction domains, including the N-terminal located LISH and CTLH domain and two blocks of WD40 repeats that are predicted to form a clamp-like structure (SZEMENYEI *et al*. 2008; KE *et al*. 2015; MARTIN-AREVALILLO *et al*. 2017). Both mutations truncated the first and second block of WD40 repeats, respectively (Supplemental Fig. 2). In Liu et al. 2019 (LIU *et al*. 2019), we reported that *RELK1* was upregulated in *rel2-ref* immature tassels while the expression levels of the other remaining family members, *RELK2* and *RELK3*, remained unchanged. RELK2 and RELK3 are phylogenetically related to AtTPL (LIU *et al*. 2019). These results suggest that an active compensation mechanism exists specifically between *REL2* and *RELK1* and agrees with the enhancement of the *rel2* phenotype in *sup* mutants. We confirmed by qRT-PCR that the increased levels of expression of *RELK1* were also observed in young vegetative tissue of *rel2-ref* mutants (Supplemental Fig. 2). From here thereafter we will refer to *sup* mutants as *rel2;relk1* double mutants.

The *RELK1* gene is expressed in every maize tissue (LIU *et al*. 2019) and we verified its expression pattern by *in situ* hybridizations in immature ears (Supplemental Fig. 2). A broad expression pattern of *RELK1* appeared in every cell similar to what was originally reported for *REL2* (GALLAVOTTI *et al*. 2010).

### Triple *rel2;relk2;relk3* mutants fail to maintain vegetative development

As mentioned above, *RELK2* and *RELK3* are two paralogous gene that belong to the TPL clade (LIU *et al*. 2019).To investigate their function, we used CRISPR-Cas9 to induce mutations in both genes, which share 94% sequence identity, with one gRNA targeting their second exon. We functionally characterized a combination containing a 16bp deletion in both genes (Fig. 1c). Double *relk2;relk3* mutant, as well as single *relk1, relk2, and relk3* mutants did not show any major developmental defects (Supplemental Fig. 3 and Supplemental Fig. 4). However, when crossed to either the *rel2-203* or *rel2-ref* allele triple mutant plants were severely impaired in shoot development (Fig. 1d; Supplemental Fig. 4). In the majority of progenies of +/*rel2;relk2;relk3* self-pollinated plants, we observed a skewed 1:1 segregation for *relk2;relk3* and *+/rel2;relk2;relk3* (Supplemental Fig. 4) indicating that most triple mutant gametes did not transmit regularly through generations. Occasionally, some *rel2;relk2;relk3* plants developed but the majority of plants failed to develop seedling leaves or when formed appeared as fused and tubular structures (Fig. 1d). Only a few rare plants made it to the reproductive phase, but those plants carried tassels with no flowers and no ears were formed (Supplemental Fig. 4). Crosses of additional alleles of *relk2* and *relk3* with *rel2-203* or *rel2-ref* recapitulated these phenotypes (Supplemental Fig. 4) that are reminiscent of the original temperature-sensitive *tpl1-1* allele of Arabidopsis (LONG *et al*. 2006). Indeed, when sectioned, these plants show a lack of formation and maintenance of the SAM. Relative to the A619 SAM, the triple mutant SAM appears to be missing (Fig. 1e). Correspondingly, we verified loss of meristem identity by *in situ* hybridizations with the meristem marker *KNOTTED1* (*KN1*). Whereas *KN1* expression was detected in the A619 SAM, this expression was absent in the triple mutant SAM remnants (Fig. 1f). In both genotypes we detected strong expression of the control marker *ZYB15* (Fig. 1f) suggesting lack of *KN1* expression in triple mutants was not due to technical issues. We also sectioned developing embryos at three developmental time points after pollination (days after pollination, DAP) and noticed that SAM development was delayed when compared to wild type embryos (Supplemental Fig. 5). By 10 DAP, wild-type and *rel2* embryos had formed the SAM and leaf primordia while in *relk2;relk3* the SAM was still exposed with only two primordia on its flanks (Supplemental Fig. 5). In *rel2;relk2;relk3* no SAM formation occurred at this stage, and *rel2;relk2;relk3* mutants were still at the pro-embryo globular stage. *relk2;relk3* was able to recover developmentally by 21 DAP (Supplemental Fig. 5) relative to wild-type embryos. By 14 DAP (Supplemental Fig. 5) *rel2;relk2;relk3* embryos were only able to form a SAM-like structure. At 21 DAP *rel2;relk2;relk3* embryos had formed several leaf primordia, however the SAM appeared flattened suggesting that halfway through embryo development the SAM was not maintained (Supplemental Fig. 5). These zygotic developmental delays, as well as potential gametophytic effects may explain the skewed 1:1 segregation for +/+;*relk2;relk3* and *+/rel2;relk2;relk3* in the majority of progenies of +/*rel2;relk2;relk3* self-pollinated plants (Supplemental Fig. 4). In order to understand whether this phenotype was due to functional diversification or a higher order effect we generated *rel2;relk1;relk3* and *relk1;relk2;relk3* mutants which phenocopied the *rel2;relk2;relk3* termination phenotype (Supplemental Fig. 5). Overall, these results indicate that REL2, RELK1, RELK2 and RELK3 function redundantly during embryogenesis and during shoot development post-germination.

To investigate possible functional diversification and redundancy among *RELK* genes, we also generated *rel2;relk2* and *rel2;relk3* double mutants. Interestingly, unlike *rel2;relk1* mutants, *rel2;relk2* and *rel2;relk3* plants reached a height similar to wild type plants. A few differences in average internode length and internode numbers relative to *rel2* were observed: for example, we found a significant decrease in average internode length of *rel2;relk3* relative to A619, but this was compensated by the formation of an additional one or two internodes in the double mutant (Supplemental Fig. 6). *rel2;relk2* mutants produced fewer tassel branches relative to *rel2;relk3*; however there were overall non-significant differences when compared to A619 and *rel2* (Supplemental Fig. 6). While all three double mutants had similar upright tassel branches (Supplemental Fig. 6), we observed differences in ear development such that 100% of *rel2;relk3* mutants failed to produce ears, while similar to *rel2;relk1, rel2;relk2* primary ears were borne on lower internodes relative to *rel2* (Supplemental Fig. 6). These results suggest that *RELK* genes have mostly redundant yet somewhat diverse functions in maintenance of various meristem types through the plant life cycle.

### REL2 and RELK1 regulate inflorescence meristem size by silencing cell-proliferative signals from various pathways

The most striking phenotype observed in *rel2;relk1* double mutants was a strong enhancement of the size of ear inflorescence meristems (fasciation; Fig. 2a), even though ears failed to develop fully and often did not produce many seeds after fertilization (Fig. 1b). We quantified the size of the inflorescence meristems and noticed that they tended to be ∼20% larger than wild type (Fig. 2b). This enhancement of ear inflorescence meristem size was similarly observed in *rel2;relk2* ears, which were ∼35% larger than wild-type and also produced small mature ears with few viable seeds (Supplemental Fig. 6). Interestingly, expression analysis of immature *rel2;relk2* ear primordia revealed that *RELK1* was strongly upregulated while *RELK3* was severely downregulated, suggesting that *RELK1* is not only unable to buffer against loss of *RELK2* function but may repress *RELK3* (Supplemental Fig. 7). Overall, *RELK1* appears as the compensatory gene within the *REL2* family as its expression was upregulated in vegetative tissue of *rel2, relk1, relk2,* and *relk3* single mutants (Supplemental Fig. 2).

**Fig. 2.**
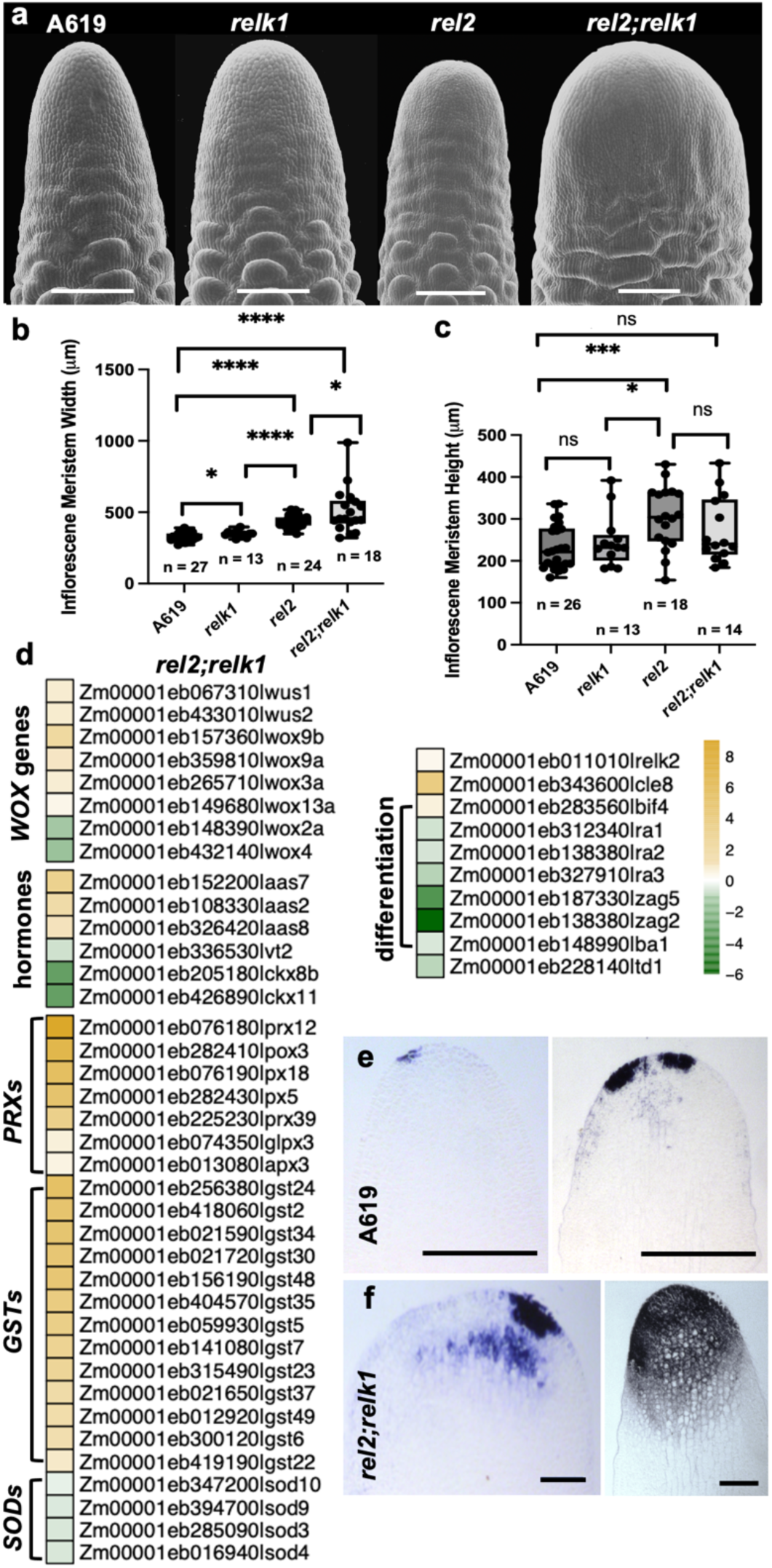
rel2;relk1 ears are fasciated. (**a**) SEM images of A619, *relk1*, *rel2*, and *rel2;relk1* ear primordia. *rel2* ears have an increase in width (**b**) leading to a cylindrically shaped IM, which becomes enhanced into a tongue shape in *rel2;relk1.* (**c**) *rel2* and *rel2;relk1* mutants do not affect IM height. (**d**) Heatmap of *rel2;relk1* differentially expressed genes in RNA-seq data (scale = LogFC) revealed mis-regulation of peroxidases (*PRXs*) and superoxide dismutases (*SODs*) which suggested an IM environment locally high in superoxide, which was confirmed by stronger NBT blue staining of (**f**) *rel2;relk1* relative to (**e**) A619 ear primordia. n = 5 per genotype. Quantification by two-tailed Student’s t-test. Box plot center line corresponds to median; box limits, upper and lower quantiles; whiskers, maximum and minimum values; ns = p > 0.05, * = p ≤ 0.05, ** = p ≤ 0.01, *** p ≤ 0.001, **** p ≤ 0.0001. Scale bars = 200μm.

To understand whether *REL2* and *RELK1* affect meristem size by influencing the core CLV-WUS pathway, we collected the first 1mm tip of immature ears in three different genotypes, A619, *rel2-ref/rel2-ref,* and *rel2-ref/rel2-ref;relk1-826/relk1-826* mutants and profiled them by RNA-seq. Differential gene expression analysis was performed, and in *rel2* tips 1,755 genes were upregulated and 1,265 genes were downregulated while in *rel2;relk1* tips 2,585 genes were upregulated and 2,130 genes were downregulated. Between these, 889 genes were similarly upregulated and 726 genes downregulated in both mutant backgrounds (Supplemental Fig. 8). Within these common genes, there was a significant mis-regulation of several transcription factor families: SBP, ARF (exclusive upregulated), EREB, DOF, WOX, AUX/IAA (predominantly upregulated), WRKY, HB, bZIP, BHLH, and MADS (equal up-and down-regulated genes) (Supplemental Dataset 1). Interestingly, in line with genetic enhancement of IM fasciation, we found an increase in the number of genes per transcriptional factor family which were mis-regulated. For example, whereas only *WOX9b* was upregulated in *rel2* tips, 5 additional *WOX* genes were mis-regulated in *rel2 relk1* tips (Fig. 2d; Supplemental Fig. 8). Similarly, we found several upregulated genes encoding reactive oxygen species (ROS) scavenging enzymes including 5 and 7 PEROXIDASES (PRXs), and 7 and 13 GLUTATHIONE TRANSFERASES (GSTs), while 1 and 4 SUPEROXIDE DISMUTASES (SODs) were downregulated in our *rel2* and *rel2;relk1* data, respectively (Fig. 2d; Supplemental Fig. 8). In addition, significant changes in expression were detected in *CYTOKININ OXIDASE* (*CKX*), *AUXIN AMIDO SYNTHETASE* (*AAS*), and the *TRYPTOPHAN AMINOTRANSFERASE VT2* genes (Fig. 2d) (PHILLIPS *et al*. 2011), which are involved in hormone degradation, conjugative inactivation, and synthesis, respectively. In line with this, GO annotation analysis performed on the *rel2;relk1* DEGs showed that these genes were highly enriched for biological processes related to response to chemical stimulus (GO:0044238, p = 2.7e-14), response to stress (GO:0006950, p = 7.2e-7), response to hormone stimulus (GO:009725, p = 0.00067), hormone-mediated signaling pathway (GO:0009755, p = 0.008), purine nucleoside triphosphate catabolic process (GO:0009146, p = 0.00027), cellular amino acid and derivative metabolic process (GO:006519, p = 1.3e-5), and auxin mediated signaling pathway (GO:0009734, p = 0.00022) (Supplemental Fig. 8). Additionally, genes related to developmental process (GO:0032502, p = 4.1e-7) and anatomical structure development (GO:0048856, p = 7.1e-6) were especially enriched and corresponded to strong down-regulation of differentiation and determination-related genes responsible for different axillary meristem identities such as *RAMOSA1* (*RA1*), *RA2*, *RA3, ZAG5, ZAG2* and *BARREN STALK1* (MENA *et al*. 1995; GALLAVOTTI *et al*. 2004; VOLLBRECHT *et al*. 2005; BORTIRI *et al*. 2006; SATOH-NAGASAWA *et al*. 2006) (Fig. 2d). In order to confirm the biological relevance of mis-regulated ROS scavenging genes we performed NBT staining for superoxide on IM tips of A619 and *rel2;relk1* (Fig. 2e-f). We detected consistently more and darker blue staining in *rel2;relk1* tips relative to A619 corresponding to enrichment in superoxide concentration, suggesting a local scrubbing of hydrogen peroxide and a concomitant increase in superoxide concentration in double mutant tips.

We also compared these results with datasets obtained in *Bif3*, a dominant mutant with enlarged ear IMs, caused by overexpression of the maize *ZmWUS1* gene (CHEN *et al*. 2021). Only 295 DEGs (differentially expressed genes) were shared between *Bif3* and *rel2* and included genes involved in various biological processes without a discernible pattern pointing to a shared regulatory mechanism (Supplemental Dataset 2). A similar range of various processes characterized the 446 DEGs which were shared between *Bif3* and *rel2;relk1*, and 3 genes involved in axillary meristem function (*RA2, RA3*, and *BARREN STALK1*) (GALLAVOTTI *et al*. 2004; BORTIRI *et al*. 2006; SATOH-NAGASAWA *et al*. 2006) were downregulated in both datasets. In a comparison of *Bif3* down-*rel2* up regulated genes one notable gene was *RELK1* suggesting it is under negative regulation by *ZmWUS1.* We also compared these results with datasets obtained in *fea4,* a semi-dwarfed mutant with fasciated ears cause by mutations in a bZIP transcription factor (PAUTLER *et al*. 2015) to see if there was overlap in DEGs between *rel2* and *fea4*. 459 DEGs were shared between *fea4* and *rel2*. In a comparison of *fea4* up*-rel2* up regulated genes, auxin related genes such as *AUX/IAAs*, and *ARFs* were enriched and this enrichment was shared in comparisons with *rel2;relk1* data (Supplemental Dataset 2). Interestingly, in a comparison of *fea4* down-*rel2* up regulated genes, a notable gene was *RELK1*, suggesting that *RELK1* is under negative regulation by *ZmWUS1* but positive regulation by *FEA4* (log2 FC -0.24).

To better understand the interaction of REL2/RELK corepressors with known meristem regulators, including core components of the CLV-WUS pathway, we generated *rel2;td1*, *rel2;fea4, rel2;wus1,* and *rel2;fea3* double mutants and *rel2;relk1;wus1* triple mutants. SEM analysis of immature ear primordia revealed synergistic effects between *rel2*, and *fea3* and *fea4* mutants (Fig. 3a-d). In the case of *rel2;fea3* double mutants, enlargement of the IM resulted in IM splitting which was not detected in *fea3* single mutants (Fig. 3d) and resulted in different mature ear morphologies (Supplemental Fig. 9). In *fea4* mutants, the IM normally splits (Fig. 3c), whereas in *rel2;fea4* we observed an enhancement in the degree of splitting. Whereas *fea4* mutants can form a mature ear with abundant seeds, *rel2;fea4* ears were smaller and failed to produce seeds (Supplemental Fig. 9). Our analysis of *rel2;td1* developing primordia and mature ears found no discernible differences between *td1* and *rel2;td1* (Fig. 3b; Supplemental Fig. 9). Occasionally, the cleft seen in the top view SEM of the ear primordia appeared wider in *rel2;td1* relative to *td1* however mature ears lack any clear differences (Supplemental Fig. 9). Overall, these data suggest an epistatic relationship between *TD1* and *REL2*, which may be in part explained by the known direct physical interaction of TPL corepressors with AtWUS. We confirmed that ZmWUS1 can physically interact with REL2 and RELK1 in Y2H assays (Supplemental Fig. 9), and may be therefore responsible for mediating ZmWUS1’s repressive activity, similar to AtTPL (CAUSIER *et al*. 2012). Therefore, taking advantage of previously generated CRISPR-Cas9 alleles of *ZmWUS1* (CHEN *et al*. 2021), we also obtained *rel2;wus1* and *rel2;relk1;wus1* mutants. We found non-significant differences in IM size between A619 and *wus1* single mutants bearing an 8bp deletion near the end of the DNA binding domain. In *rel2;wus1* and *rel2;relk1;wus1* we observed a significant rescue of fasciation (Fig. 3a,e) in which the morphology of the meristem returned to a normal conical shape, but width was still significantly increased relative to A619. Take together with the upregulation of *ZmWUS1* (Fig. 2c), our data suggests that *rel2;relk1* mutants promote a molecular environment which tips the meristem toward fasciation by expanding *ZmWUS1* expression. The absence of a complete rescue by the loss of function mutation in *ZmWUS1* indicates that other stem-cell promoting factors contribute to the fasciated phenotype, in accordance with the differential expression analysis.

**Fig. 3.**
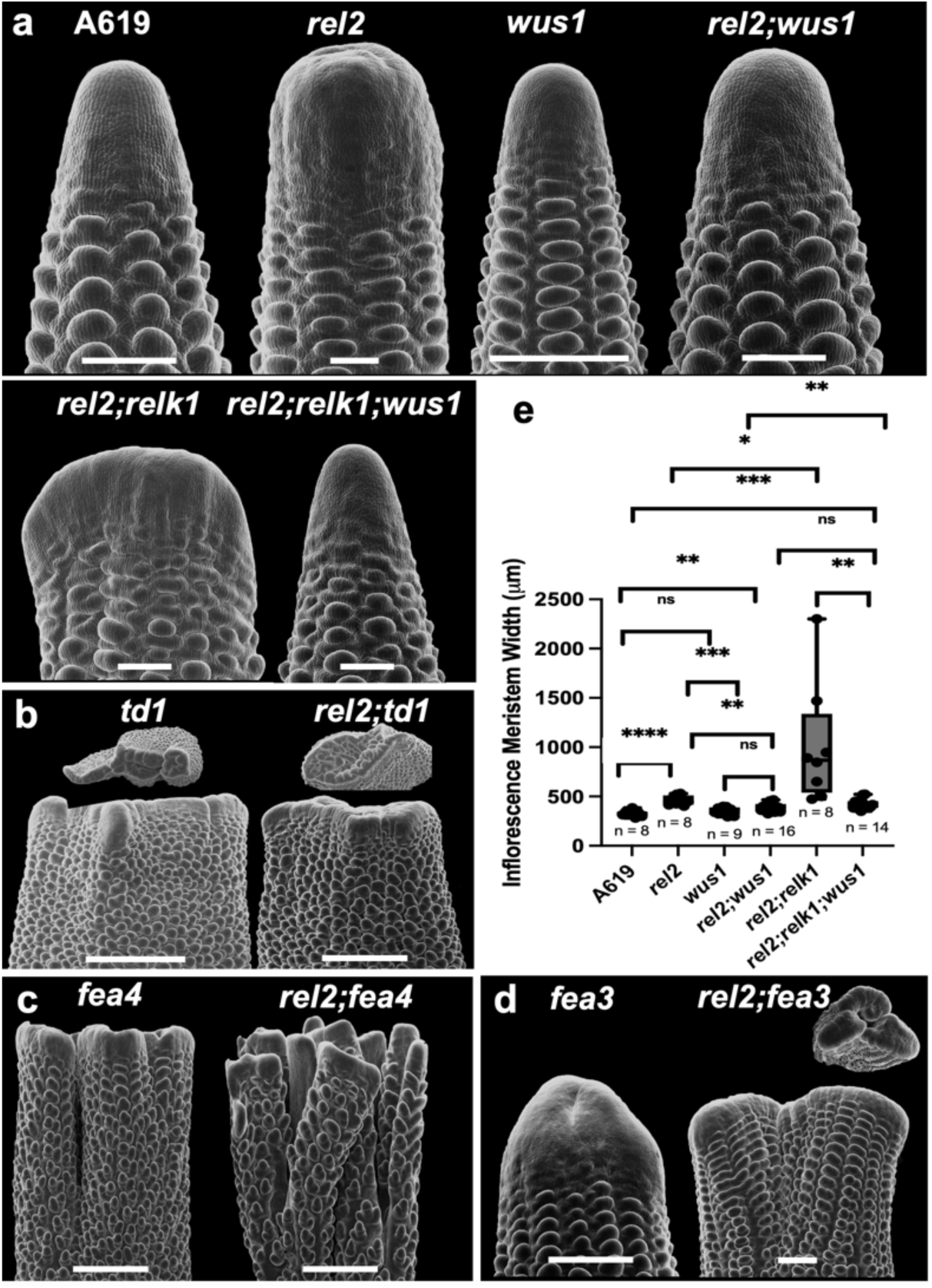
Genetic interaction analysis with fasciated mutants. (**a,e**) Introduction of a CRISPR-Cas9 generated *wus1* deletion allele into the *rel2* and *rel2;relk1* backgrounds rescued IM morphology and tempered but did not entirely rescue IM width relative to A619. Recovery of a conical IM in *rel2;relk1;wus1* suggests all three operate within the same pathway. (**a**) Relative to *rel2*, (**b**) *td1* and *rel2;td1* barrel-shaped ear primordia show no discernible differences in line with an epistatic interaction. Whereas the presence of *rel2* in the (**c**) *fea4* and (**d**) *fea3* background resulted in strong synergistic splitting of the IM relative to each single mutant. A619, *wus1* scale bar = 200μm; *rel2, fea4, fea3, rel2;fea3, rel2;fea4, rel2;wus1* scale bar = 500μm; *td1, rel2;td1* scale bar = 1mm. Quantification by two-tailed Student’s t-test. Box plot center line corresponds to median; box limits, upper and lower quantiles; whiskers, maximum and minimum values; ns = p > 0.05, * = p ≤ 0.05, ** = p ≤ 0.01, *** p ≤ 0.001, **** p ≤ 0.0001. SEM images, n = 8-16 per genotype.

### Mutations in *REL2* increase seed yield

We previously reported that single *rel2* mutants in the permissive A619 background produced subtle increases in ear IM size and subsequently mature ears with an increase in rows of kernels, a yield trait called KRN (Kernel Row Number) (LIU *et al*. 2019). In B73 however, *rel2-ref* homozygous mutants failed to develop ears (LIU *et al*. 2019). We subsequently noticed a similar subtle increase in IM size in *rel2-ref* heterozygous plants in the B73 inbred background (Fig. 4a-b) that produced ears with a tendency to form two extra rows of kernels (Fig. 4c). This heterozygous effect on ear size prompted us to test it in F1 hybrids, which are commonly used for commercial seed production. We first tested the *rel2* heterozygous effect in F1 hybrids obtained by crossing A619 (+/+) and B73 (*rel2-ref/rel2-ref*). Results from three independent fields indicated that +/*rel2* F1 hybrids produced two extra rows of kernels on average (p<0.0001) when compared to A619 (+/+) X B73 (+/+) hybrids (Fig. 4d,g). The size of the ears and kernel weight were not significantly affected by the *rel2* mutation, apart for a subtle increase in length (Fig. 4e-f; Supplemental Dataset 3). To verify whether this effect was widespread among different hybrid combinations and heterotic groups, we also generated 36 additional F1 hybrids, using as parents mostly a subset of the NAM founder lines, which capture different maize groups and a majority of variability existing in maize germplasm, as well as other available inbreds (Supplemental Dataset 3) (YU *et al*. 2008). We carried out limited field trials in at least two independent fields, either in different years or in different locations (NJ and HI) for additional 22 of the 36 F1 hybrids. Overall, 40% of the tested hybrids with replicates produced significant increases in KRN (Fig 4j; Supplemental Fig. 10); however, only in 4 additional combinations (OH43xB73, CML69xA619, M37WxB73 and Ms71xB73) were these increases observed without adversely affecting ear size (Fig. 4h) or kernel weight (Fig. 4i), suggesting that heterozygosity at the *rel2* locus can potentially be used to significantly increase maize seed yield in a subset of hybrid combinations.

**Fig. 4.**
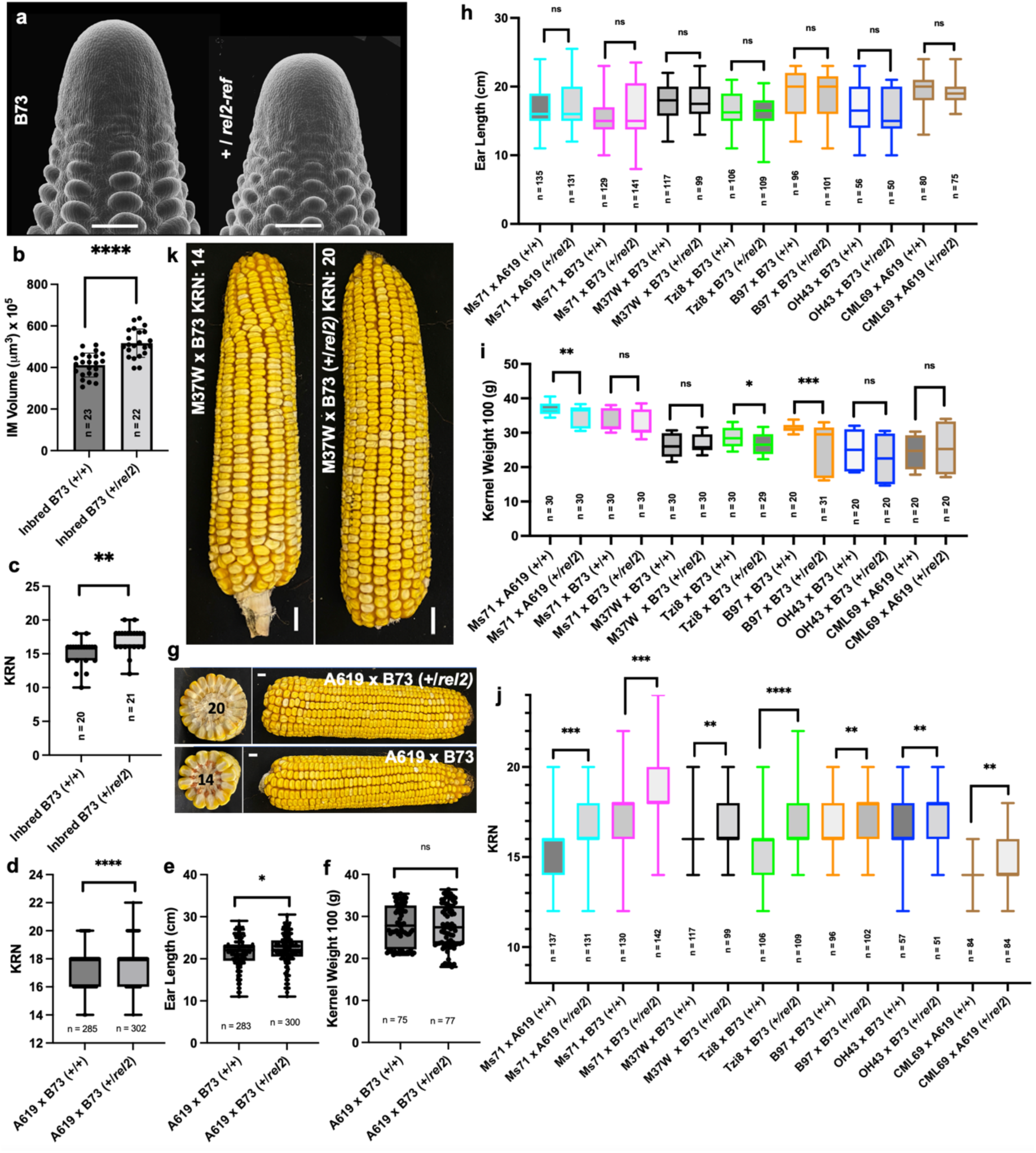
The *rel2-ref* allele increases KRN. (**a**) SEM images of B73 wild-type and +/*rel2* ear primordia. Scale bars = 200μm. The presence of a single mutant allele results in a significant increase in IM volume (**b**). Concomitantly, heterozygous mutant inbred lines (**c**) and (**d-g**) A619 x B73 F1’s display an increase in KRN relative to wild type controls without affecting kernel weight; only a subtle increase in length was observed in +/*rel2* hybrids. (**h-j**) 22 additional F1 hybrid combinations were tested for increased KRN. Of these, (**j**) 7/22 displayed increased KRN in 2-3 environmental replicates without significant alterations in ear length (**h**), though pleiotropic effects on kernel weight (**i**) were observed. (**k**) Representative image of M37W x B73 (+/+) and M37W x B73 (+/*rel2*) ears showing an increase KRN without morphological deformity of the ear. Scale bars = 1 cm. Quantification by two-tailed Student’s t-test. Box plot center line corresponds to median; box limits, upper and lower quantiles; whiskers, maximum and minimum values; ns = p > 0.05, * = p ≤ 0.05, ** = p ≤ 0.01, *** p ≤ 0.001, **** p ≤ 0.0001.

## DISCUSSION

Transcriptional corepressors are master regulators of development throughout the Eukarya (LONG *et al*. 2006; PAYANKAULAM *et al*. 2010; TURKI-JUDEH AND COUREY 2012; BAILEY *et al*. 2022). Previous studies from Arabidopsis, rice and maize have highlighted their importance within the Angiosperms in regulating developmental programs and immune responses (LONG *et al*. 2006; GALLAVOTTI *et al*. 2010; PAUWELS *et al*. 2010; ZHU *et al*. 2010; YOSHIDA *et al*. 2012; WALLEY *et al*. 2018; LIU *et al*. 2019; SUZUKI *et al*. 2019a; DARINO *et al*. 2021; PLANT *et al*. 2021; HUANG *et al*. 2024). In this study, we provide a comprehensive functional analysis of the *REL2* corepressor family members and their indispensable role in maize growth. Double and triple mutant analysis of family members revealed redundant and divergent roles of individual *RELK* genes. The SAM termination and early embryogenic developmental delays in *rel2;relk2;relk3* and *relk1;relk2;relk3* indicate that REL2, RELK1, RELK2, and RELK3 function redundantly during embryogenesis and shoot development post-germination, in a similar fashion to the Arabidopsis *tpl1-1* allele, a gain-of-function dominant negative mutant (LONG *et al*. 2006). We detected a skewed 1:1 segregation for *relk2;relk3* and *+/rel2;relk2;relk3* in the majority of progenies of +/*rel2;relk2;relk3* self-pollinated plants which may be partially explained by developmental delays during early embryogenesis, or potential gametophytic defects; however further investigation is required to parse out whether one or both causes contribute to the observed segregation. With the switch to the reproductive phase, we have shown that *REL2, RELK1* and *RELK2* are required for maintenance of ear IM size, while *REL2* and *RELK3* function in the initiation of the axillary buds which will form ears. Taken together, our data suggests that *REL2, RELK1, RELK2,* and *RELK3* are necessary for the initial establishment and maintenance of the embryonic SAM, but *REL2* functions as the essential gene, while the other members appear dispensable and can only partially compensate for its loss. Given the phenotypic consequences of single or higher order mutations among a small group of vital proteins, evolution would favor redundancy within the system to buffer against lethality and loss of reproductive fitness.

The most intriguing phenotype was the enhanced fasciation in *rel2;relk1* relative to *rel2* ears. Understanding the mechanism of fasciation has important implications for fine-tuning yield increases. Based on our analysis, *REL2* and *RELK1* act cooperatively in the IM to repress cell proliferation in the meristem thereby ensuring homeostasis and regulating size by controlling genes involved in redox balance, hormone catabolism, and differentiation. This includes the up-regulation of several AASs and a concomitant down-regulation of CKXs and VT2 (PHILLIPS *et al*. 2011). This should result in an environment low in free, biologically available auxin but high in cytokinin, an essential hormone promoting stem cell proliferation. Prior work with the *barren inflorescence* mutants has demonstrated the importance of both hormones in maintaining meristems or promoting differentiation. The semidominant *Bif3* (CHEN *et al*. 2021) mutant causes stem cell over-proliferation in the IM due to changes in cytokinin sensitivity at the *ZmWUS1* locus, while studies of *bif2* (MCSTEEN *et al*. 2007) and *Bif1, Bif4* (GALLI *et al*. 2015) have demonstrated the vital role of auxin in establishing axillary meristems as the ear develops. From a redox standpoint, some of the most numerous and highest differentially expressed genes corresponded to *PRXs* and *GSTs* which detoxify hydrogen peroxide (UGALDE *et al*. 2021). Together with the concomitant downregulation of *SODs*, the *rel2;relk1* female IM should be scrubbed of differentiation promoting hydrogen peroxide and locally high in the stem-cell promoting superoxide, a hypothesis that was supported by the NBT staining assay (Fig. 2). Superoxide can be protonated to the reactive perhydroxyl radical which can initiate lipid peroxidation chains (DAT *et al*. 2000). GSTs function in the detoxification of hydrogen peroxide and lipid peroxides, with different enzymes bearing different affinity for one or the other (UGALDE *et al*. 2021). Therefore, the large number of upregulated *GSTs* may reflect not only additional means to mute differentiation but also tamper damage to cellular ultrastructure in the presence of high superoxide conditions.

Our genetic analysis between *rel2* and already characterized fasciated mutants indicates that *REL2/RELKs* mainly function in the CLV-WUS pathway, but separately from both *FEA3* and *FEA4* pathways (Fig. 3). Based on the interaction analysis of TPL and WUS in Arabidopsis, this is expected, and WUS, a bifunctional transcription factor (IKEDA *et al*. 2009), is believed to work as a repressor in the organizing center via its physical interaction with TPL proteins and repress genes involved in differentiation (KIEFFER *et al*. 2006; CAUSIER *et al*. 2012). However, this simple model does not explain why loss-of-function mutations in *REL2* (and *RELK* genes) and its rice ortholog *ASP1* increase inflorescence meristem size, rather than decreasing it, as predicted. In rice, mutations in *TAB1*, the *WUS* ortholog, do not affect meristem size but rather the initiation of axillary meristems (SUZUKI *et al*. 2019a; SUZUKI *et al*. 2019b) while in maize loss of *ZmWUS1* activity does not affect development (CHEN *et al*. 2021), presumably due to the presence of the duplicated gene *ZmWUS2* (NARDMANN AND WERR 2006). However, by generating *rel2;wus1* and *rel2;relk1;wus1* mutants we clearly showed an almost complete rescue of the fasciated ear phenotype, indicating that *ZmWUS1* overexpression observed in our RNA-seq datasets is largely responsible for the increase in meristem size. Rather than simply acting through a physical interaction with WUS proteins, our data supports a broader role for REL2-RELK in maintaining meristem homeostasis by influencing the CLV-WUS pathway, the auxin/cytokinin balance and ROS species. Overall, these factors maintain a molecular environment that tampers *ZmWUS1* expression, whose repressive activity may require REL2/RELKs.

In the same way the *REL2* family is indispensable to maize, maize is an important direct or indirect source of food. As 30% of the world’s caloric intake can be traced directly or indirectly back to maize (SHIFERAW *et al*. 2011), even modest increases in seed yield per acre on a commercial scale can have large impacts on maintaining vital food chains. Through our hybrid analysis, we have shown the potential application of *rel2* mutants in significantly increasing maize yield. While only 20% of our F1 hybrid combinations resulted in a significant increase in KRN without detrimental pleiotropic effects, it is possible that our analysis failed to capture all potential additive variance by using only two inbred lines as sources for the *rel2-ref* allele. While these results are promising, we caution that large-scale production tests should be performed to thoroughly evaluate the yield performance of these and other hybrid combinations (KHAIPHO-BURCH *et al*. 2023). Also, REL2 has been implicated in resistance to biotic stresses (WALLEY *et al*. 2018; DARINO *et al*. 2021; BINDICS *et al*. 2022; HUANG *et al*. 2024); whether *rel2* heterozygous plants are more susceptible to pathogens or other stresses remains to be properly tested.

## MATERIALS AND METHODS

### Plant Material and Positional Cloning of *RELK1*

Maize plants were grown in the Waksman Institute field (Piscataway, NJ, USA) or in Rutgers Hort Farm III (East Brunswick, NJ, USA) during the summer months, or during the winter in the Waksman Institute greenhouse (Piscataway, NJ, USA) or in the Friendly Isle Growing Service Corporation field nursery in Hawaii (Molokai, HI, USA).

The *relk1-826* and *relk1-913* alleles were generated by EMS mutagenesis in a screen for enhancers of the *rel2-ref* mutant phenotype (LIU *et al*. 2021). The original mutagenesis was performed in a permissive background (A619). Both original *rel2;relk1* double mutants were then crossed to B73 to generate F2 mapping populations. A whole-genome sequencing bulked segregant analysis was subsequently performed by sequencing DNA bulk samples from pools of double mutant plants (14 *rel2;relk1-826* and 18 *rel2;relk1-913* plants). Library preparation and sequencing were performed by Macrogen/Psomagen using Illumina HiSeq X Ten and NoveSeq 6000 S4 systems, with an approximate 15x and 50x coverage, for *rel2;relk1-913* and *rel2;relk1-826* respectively. Sequence analysis and SNP calling were done according to Dong et al. 2019 (DONG *et al*. 2019).

To obtain CRISPR-Cas9 mutant alleles of *RELK2* and *RELK3* a dual targeting guide RNA specific for exon 2 was cloned into pBUE411 by HiFi cloning (NEB). To obtain CRISPR-Cas9 mutant alleles of *RELK1* one guide RNA specific for exon 2 was cloned into pBUE411 by HiFi cloning (NEB). The resulting construct was introduced into Hi-II immature embryos by Agrobacterium-mediated transformation, followed by callus induction, bialaphos selection, and shoot and root generation. Transformation and regeneration were carried out by the Iowa State University Plant Transformation Facility. Mutant alleles were generated by crossing transgenic lines carrying CRISPR-Cas9 *RELK1;RELK2;RELK3* gRNA with A619 wild-type, *rel2-ref;relk1-826* and *rel2-203* mutant (A619) BC(2). Genotypes for single, double, and triple mutants were determined using gene-specific primers (Supplemental Table 1). For the genetic analysis, we generated double mutants by crossing *td1-mum3*, *fea4-5171, fea3-0* alleles with either *rel2-ref* (A619) or *rel2-203* (A619) (BOMMERT *et al*. 2005; PAUTLER *et al*. 2015; JE *et al*. 2016; LIU *et al*. 2019). For quantification two additional rounds of back-crosses in A619 were performed. To generate *rel2;wus1* mutants, we crossed CRISPR-Cas9 *ZmWUS1*-A specific guide RNA line (CHEN *et al*. 2021) with *rel2-ref;relk1-826*. Quantification was carried out after three rounds of back-crosses in A619.

Field-grown *rel2-ref;relk1-826* double mutants A619 BC(6) were phenotyped for plant height (aerial root internode to tassel tip), internode number, internode length, internode at which the primary ear was formed, length of terminal internode to tassel tip, length of subtending tassel branch to tassel tip, and tassel branch number in summer 2020 and 2021 field seasons. Measurements included at least 10 samples per genotype, with significance calculated using student’s two-tailed *t-*test, with a p value <0.05 threshold. Graphpad Prism 9 software was used to represent the data.

### Scanning Electron Microscopy

Freshly dissected samples of immature ears (3 mm – 5mm) were imaged using a JMC-6000PLUS Scanning Electron Microscope within 15 minutes of dissection. IM width and height measurements were taken immediately after images were captured. All measurements included at least 8 samples per genotype, with significance calculated using student’s two-tailed *t-*test, with a p value <0.05 threshold. Graphpad Prism 9 software was used to represent the data.

### Sectioning and Histology

For analysis of SAM termination, A619 wild-type and *rel2-203;relk2;relk3* seedling tissue was harvested from samples grown in the Waksman Institute greenhouse in the spring of 2021. For analysis of early embryogenesis 10 DAP, 14 DAP, 16 DAP and 21 DAP immature kernels were harvested from self-pollinations of A619 wild-type, *rel2-203, relk2;relk3,* and +/*rel2-203;relk2;relk3* ears grown in the Waksman Institute field in the summer of 2021. Immature kernels were genotyped using endosperm tissue with *rel2-203* primers (18; Supplemental Table 1). Tissues were fixed in 4% PFA (paraformaldehyde; Thermo Scientific) using vacuum infiltration, then dehydrated in a graded ethanol series, treated with a graded Histoclear series (Electron Microscopy Solutions), and embedded in paraplast (Leica).

Sections (8μm) were placed on slides (Fisherbrand), dewaxed, and stained with 0.1% Toluidine Blue (in 0.6% boric acid). Stained sections were rinsed with water, mounted with Permount (Electron Microscopy Solutions), and analyzed by light microscopy using a Leica DM5500B microscope equipped with a DFC450 C digital camera.

### NBT Staining

Nitro blue tetrazolium staining was performed on 2-5 mm ear primordia harvested from A619 and *rel2;relk1*. Primordia were incubated in 1mg/ml NBT solution (50 mg NBT in 50 mL 6.8g/l KH2PO4 with 0.05% v/v Tween 20) under vacuum for 3 hours and incubated overnight in the dark. Following incubation, the NBT solution was replaced with 3:1:1 ethanol:acetic acid:glycerol solution and boiled at 95°C for 15 minutes. Then fixed in PFA (Thermo Scientific), dehydrated in a graded ethanol series, cleared with Histoclear (Electron Microscopy Solutions) and embedded in Paraplast (Leica). Sections (8μm) were placed on slides, and dewaxed. Stained sections were rinsed with water, mounted with Permount (Electron Microscopy Solutions), and analyzed by light microscopy using a Leica DM5500B microscope equipped with a DFC450 C digital camera.

### RNA In Situ Hybridizations

In situ hybridization experiments were performed on 2 - 5 mm ear primordia and 10-21 day old seedling apexes fixed in PFA (Thermo Scientific), dehydrated in a graded ethanol series, cleared with Histoclear (Electron Microscopy Solutions) and embedded in Paraplast (Leica). Sense and antisense RNA probes for *ZmRELK1*, *ZYB15,* and *KN1* genes, using either entire/partial coding sequences or 3′-untranslated region of each gene were cloned into pGEM T-easy vectors (Promega). The plasmids were linearized by restriction endonucleases, and the sense/ antisense RNA probes (with sizes ranging from 300 to 1000 bp) were synthesized by T7 or SP6 RNA polymerase (Promega). The vectors, enzymes, and primers used for probe design are listed in Supplementary Table 1.

### RNA-seq and Expression Analysis

Immature ears (3 mm – 5mm) were collected from A619 wild-type, *rel2-ref* and *rel2-ref;relk1* plants grown in winter 2021 and 2022 in the Waksman Institute greenhouse. Inflorescence meristems, defined as 1 mm of tissue at the tip of immature ears, were harvested per genotype per biological replicate (a total of 20-60 tips) approximately at the same time of the day (from 11am-2pm). Total RNA was isolated using Plant RNeasy Mini Kit (Qiagen). Three biological replicates per genotype were sent to Psomagen Inc. for library preparation and Illumina RNA sequencing. RNA-seq raw data were trimmed with Trimmomatic with default settings (BOLGER *et al*. 2014). Alignment to the reference genome (B73v5 genome) was performed using HISAT 2.1.0 with default settings (KIM *et al*. 2015). HTSeq-count (ANDERS *et al*. 2015) was used to quantify gene expression as read counts and genes with differential expression were determined by edgeR 3.18.1 package (CHEN *et al*. 2016). All statistical analyses of gene expression were conducted in R. Genes with a fold change *p* < 0.05 were considered as differentially expressed genes. GO Term Analysis was performed using AgriGO v2.0 GO Analysis Toolkit and Database. All RNA-seq datasets are deposited in NCBI (BioSample accession numbers SAMN40984045-SAMN40984074).

For RT-qPCR female inflorescence analysis, immature ears (3mm - 5mm) were collected from A619 wild-type, *rel2-ref, rel2;relk2,* and *rel2-ref;relk1* plants. Total RNA was extracted from three biological replicates with 9 ears per replicate using Plant RNeasy Mini Kit (Qiagen). For the vegetative analysis, 1 centimeter of tissue (hypocotyl to leaf sheaths) were harvested from two-week old seedlings. Total RNA was extracted from three biological replicates with 3 seedlings per replicated using Plant RNeasy Mini Kit (Qiagen). cDNA was obtained using the qScript cDNA Synthesis kit (Quantabio) and amplified with the Perfecta SYBR Green Fast Mix (Quantabio). RT-qPCR was performed on two biological replicates with three technical replicates using specific primer pairs for REL2 and RELK genes ((LIU *et al*. 2019); Supplemental Table 1), and UBIQUITIN as a control. Reactions were carried out using the Illumina Eco Real-Time PCR System and quantified with the Eco Real-Time PCR System Software EcoStudy (Illumina).

### Yeast 2-hybrid

To generate the pDEST-DB-REL2, pDEST-DB-RELK1, pDEST-DB-RELK2, pDEST-DB-RELK3, and pDEST-AD-WUS1 constructs, the full-length coding sequences of REL2, RELK1, RELK2, RELK3, and WUS1 were cloned into pENTR223-Sfi and recombined into either pDEST-DB or pDEST-AD using LR clonase II (Life Technologies). pDEST-AD and pDEST-DB clones were transformed into mating compatible yeast strains Y8800 and Y8930, respectively, using the LiAc transformation method. Mating was carried out according to standard procedures. Reporter gene activation was determined by assessing growth on -Leu/-Trp/-His +1 mM 3AT media after 3 to 5 days grown at 28°C to 30°C.

### F1 Hybrid Analysis

The analysis of the *rel2-ref* allele with kernel row number (KRN) was conducted with a maize panel of 23 diverse inbred lines. F1 hybrids resulting from crosses of wild-type and *rel2-ref* mutant inbred lines in the A619 and B73 backgrounds with the 22 diverse inbred lines were grown in three environments: Molokai (HI), Waksman Institute field (WIM), and Rutgers University Horticulture Farm III (HF). Crosses were performed using the *rel2-ref* mutant as female or male for A619 hybrids, and male only for B73 hybrids (*rel2-ref* is earless in B73 (LIU *et al*. 2019)). KRN, ear length, and weight of 100 kernels phenotypic data was collected from plants grown between winter 2020 and winter 2022 (raw data are available in Supplemental Dataset 3). Weight of 100 kernels was used as a proxy for seed size. Statistical analysis was performed using Student’s two-tailed *t-* test, with a p-value <0.05 threshold, using data from all replicates. Graphpad Prism 9 software was used to represent the data. Please note that in several instances the number of ears analyzed for KRN or ear length do not necessarily match; i.e. this was caused by the fact that ear length could be measured but not KRN, due to poor seed set; or vice versa, KRN could be measured but not ear length because the ear broke off.

## AUTHOR CONTRIBUTIONS

J.G., X.L., Z.C., M.G., A.G. designed research and analyzed data; J.G., X.L., Z.C., C.G., J.P., G.K., A.G. performed research; J.G. and A.G. wrote the paper; A.G. secured funding.

## ACKNOWLEDGMENTS

We are grateful to Tim Chionis and John Bombardiere for greenhouse and field management; to the Advanced Plant Genetics class of the 2022 fall semester for help in collecting KRN data; to Fang Yang for sharing the NBT staining protocol; to David Jackson and Ayushi Agrawal for helpful discussions. This work was supported by grants from the US National Science Foundation to A.G. (IOS-1456950 and IOS-2026561).

J.G. was also supported by the Charles and Johanna Busch Graduate Fellowship in Molecular Biology at the Waksman Institute of Microbiology.

